# Semiconductor augumented valuable chemical photosynthesis from *Rhodospirillum rubrum* and mechanism study

**DOI:** 10.1101/2023.04.11.532515

**Authors:** Lin Wang, Shulan Shi, Jun Liang, Bo Wang, Xiwen Xing, Cuiping Zeng

## Abstract

Photosynthetic biohybrid systems based on purple bacteria and semiconducting nanomaterials are promising platforms for sustainable solar-powered chemical production. However, these types of biohybrid systems have not been fully developed to date, and their energy utilization and electron transfer mechanisms are not well understood. Herein, a *Rhodospirillum rubrum*-CdS biohybrid system was successfully constructed. The photosynthetic activity and photoelectrochemical properties of biohybrid system were analyzed. Chromatographic and spectroscopic studies confirmed the metabolic activities of *R. rubrum* cells were effectively augmented by surface-deposited CdS nanoparticles and validated with increased H2 evolution, polyhydroxybutyric acid (PHB) production, and solid biomass accumulation. Energy consumption and metabolic profiles of *R. rubrum*-CdS biohybrid system exhibited a growth phase-dependent behaviour. Photoelectrochemical study confirmed that light-excited electrons from CdS enhanced photosynthetic electron flow of *R. rubrum* cells. Monochromatic light modulated photoexcitation of biohybrid system was utilized to explore interfacial electron transfer between CdS and *R. rubrum* cells, and the results showed that CdS enhanced the utilization of blue light by *R. rubrum* cells. This work investigated the feasibility and prospect of utilizing *R. rubrum* in semi-artificial photosynthesis of valuable products, and offered insights into the energy utilization and the electron transfer mechanism between nanomaterials and purple bacteria.

## 1. Introduction

In recent years, scientists have discovered that semiconducting nanomaterials with light-capturing activities can assist microorganisms in converting and utilizing solar energy.^1^ The photosynthetic biohybrid systems are constructed via combining different kinds of semiconducting nanomaterials and microbial cells. As a results, several non-photosynthetic microorganisms, such as the acetogens,^2^ *Escherichia coli*^3^ and yeasts^4^ etc., were transformed into hybrid lifeforms that can carry out light-driven chemical synthesis.^5 6^ This area of study expanded people’s understanding on traditional photosynthesis, and has been termed “semi-artificial photosynthesis” in 2018.^7^ Compared to the non-photosynthetic microorganisms, photosynthetic microorganisms have the potential to uptake exogenous electrons at multiple sites of the photosynthetic electron transport chain,^8^ a property that underlines their potential compatibility with semiconducting nanomaterials. In addition, they are capable of synthesizing products from carbon dioxide, and some of them also perform nitrogen fixation.^9^ Therefore, the photosynthetic microorganisms have become a good research platform for constructing semi-artificial photosynthetic systems.

Oxygenic photosynthetic microbials, especially cyanobacteria, have received widespread attentions in semi-artificial photosynthesis studies. Zeng *et al*. designed a biohybrid system where the photoactive cationic poly fluorobenzene derivative (PFP) bind to *Synechococcus sp*. via electrostatic interaction.^10^ PFP accelerated the electron transfer in the photosynthetic electron transduction chain, which enhanced the light-dependent reactions of photosynthesis. Chen *et al*. successfully constructed a biohybrid system with molybdenum disulfide (MoS_2_) nanosheets and *Nostoc sphaeroides*,^11^ and the photoelectrons generated by MoS_2_ enhanced the production of a range of compounds from the cyanobacterial cell. However, the released O_2_ during oxygenic photosynthesis has possibility to react with the photoelectrons produced by semiconducting nanomaterials, thus hinder the electron transfer to microbial cells.^12^ Moreover, this process would also generate reactive oxygen radicals (ROS) that cause adverse effects on microorganisms.^13^ In comparison, the widely existed purple bacteria which conduct anoxygenic photosynthesis and can also produce a diverse range of high-value products for different applications,^14^ exhibited better compatibility and biosafety for constructing biohybrid systems, which was verified in a few studies.^15 16^ As a typical purple bacterium, *Rhodospirillum rubrum* has been studied as a model system for photo-to-chemical energy conversion via its nitrogen fixation pathway.^17^ It has also been used in the production of poly hydroxybutyric acid (bioplastic precursor),^18^ and generate green energy via producing hydrogen.^19^ Therefore, the investigation of *R. rubrum* for developing photosynthetic biohybrid system is promising, and the practical feasibility is worthy of exploration.

A critical factor affecting the performance of light-driven chemical synthesis by the biohybrid systems is the effective electron transfer between nanomaterials and microbial cells.^20 21^ However, the electron transfer pathways are quite different from each other between different microorganism species, and the electron transfer mechanism in biohybrid systems based on purple bacteria is still unclear. Fluorescence spectroscopy can be used to analyze the electronic structure, energy levels of molecules and carrier recombination efficiency to determine the path of electron transfer.^22^ Photoelectrochemical technology can be applied to study the electron transfer phenomenon by measuring the electrochemical reaction at the interface between microbe and material from macroscopic to microscopic levels.^23 24^ These techniques have a number of unique advantages: the wide range of accessible wavelengths (from ultraviolet to infrared), real-time and in-situ data collection, as well as the tuneable lighting conditions.^25^ Therefore, making good use of the above techniques is beneficial to analyze the unknown photogenerated electron utilization mechanism in the biohybrid system from different angles.

In this study, to explore the application prospects of *R. rubrum* in semi-artificial photosynthesis system, we constructed the biohybrid system of *R. rubrum*-CdS via a biomineralization strategy and investigated the changes in light-driven production of chemicals by bacterial cells, such as the production of hydrogen, PHB, pigments and the biomass content. We also investigated the underlying mechanisms for light-augmented metabolic activities of *R. rubrum*-CdS biohybrid system, and further explored the electron transfer at nanomaterial-cell interface with the help of spectroscopy and electrochemical techniques. This study is expected to accelerate the development and application of the sustainable chemical production by *R. rubrum*-based biohybrid systems and offer insights into the electron transfer mechanisms between semiconducting nanomaterials and purple bacteria.

## 2. Results and discussion

### 2.1 Characterization of the *R. rubrum*-CdS biohybrid

We performed biomineralization on *R. rubrum* cells using Cd(NO_3_)_2_ and *L*-cysteine hydrochloride to obtain the *R. rubrum*- CdS biohybrid system (Fig. 1a). The optimal Cd^2+^ concentration that *R. rubrum* cells could tolerate was firstly tested. After a culture period of 24 h, the OD_600_ measurements on *R. rubrum* cultures supplemented with different concentrations of Cd(NO_3_)_2_ and 1 mM of *L*-cysteine hydrochloride (Fig. 1b) gave rise to a dose-dependent inhibition pattern, and further increasing the Cd^2+^ concentration up to 0.3 mM resulted in a minor reduction in cell density than that of the 0.2 mM. Then the hydrogen production of *R. rubrum* incubated with different concentrations of cadmium ions in 24 hours was measured. It was found that the hydrogen production of *R. rubrum* incubated with 0.2 mM of Cd^2+^ was the highest of all tested Cd^2+^ concentrations (unpaired T-test, **p < 0.01) (Fig. 1c). Quantification of remaining Cd^2+^ in the supernatant of culture media indicated more than 99.9% of initially added Cd^2+^ had been converted into CdS for all tested Cd^2+^ concentrations (Fig. S1). The SEM images shown the evenly distributed nanosized bright dots on bacterial surface at Cd^2+^ concentration of 0.2 and 0.3 mM (Fig. 1d), and the low-magnification SEM image of *R*.*rubrum* treated with a Cd^2+^ concentration of 0.2 mM shown the surface of almost all cells were covered with CdS (Fig. S2). The EDS mapping on a single *R. rubrum* cell with surface-adhered bright dots (Fig. 1e) revealed the dots were composed of Cd and S elements, and the C and N elements were originated from the cell (Fig. 1f-i).

**Figure 1.**
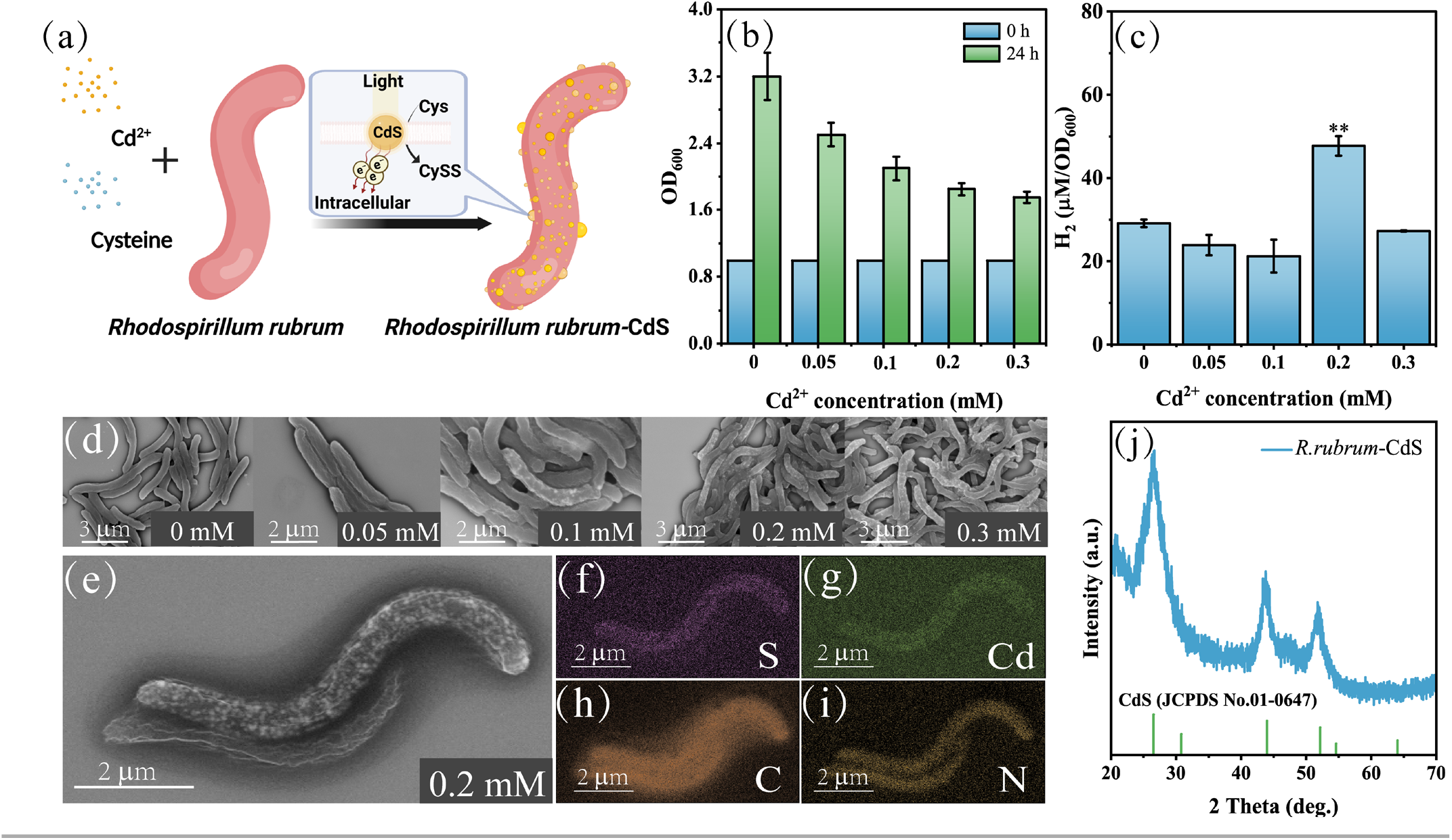
(a) schematic illustration of the formation of CdS nanoparticles on the surface of *R. rubrum* cells. (b) cell densities of R. rubrum cultured for 24 h at the presence of different concentrations of Cd2+. (c) hydrogen evolution of *R. rubrum* at different concentrations of Cd2+ in nitrogen atmosphere. (d) SEM images of *R. rubrum* cells treated with different concentrations of Cd2+. (e) SEM images of a single *R. rubrum* cell cultured under 0.2 mM of Cd2+ with surface-deposited CdS nanoparticles, and its EDS elemental maps for the elements of (f) S, (g) Cd, (h) C, and (i) N, respectively. (j) XRD patterns of *R. rubrum*-CdS biohybrid system and the reference peaks from JCPDS data card No. 01-0647 (given as drop lines)

In addition, we recorded the UV-Vis diffuse reflection spectra of the *R. rubrum*-CdS between 200 - 800 nm and converted into the Tauc plot (Fig. S3). The bandgap value of surface-deposited CdS nanoparticles, as calculated from the Tauc plot, is 2.37 eV. Powder X-ray diffraction showed broad peaks consistent with small particles of the hawleyite phase (Fig. 1j). These findings confirmed the successful deposition of CdS nanoparticles on the surface of *R. rubrum* cells, and the *R. rubrum*-CdS biohybrid system constructed with 0.2 mM Cd^2+^ was chosen for following investigations.

### 2.2 Metabolic profiles of *R. rubrum*-CdS biohybrid system

Then we evaluated the photosynthetic performance of *R. rubrum*-CdS biohybrid system via analyzing the key metabolites of the system. In purple bacteria, intracellular nitrogenase catalyses nitrogen fixation and concurrently releases hydrogen (H_2_), a process that consumes electrons delivered by ferredoxin (Fd), the electron carrier that shuttles electrons from the photosynthetic electron transport chain to various downstream enzymes (e.g., nitrogenase and NADP reductase).^26^ Since the catalytic activities of nitrogenase behaves differently under N_2_ and Ar atmosphere ^27^, the light-driven H_2_ evolution for both the native cells and the biohybrid system was evaluated under N_2_ and Ar, respectively. The biohybrid system generated more H_2_ than the native cells under identical culture conditions for both N_2_ and Ar atmosphere (Fig. 2a). The solar-hydrogen conversion efficiencies of *R. rubrum*-CdS biohybrid system and native cells were 0.102 % and 0.044 % under argon, 0.046 % and 0.025 % under nitrogen, respectively, indicating that the energy conversion efficiency of the biohybrid system was 132% higher in argon and 84% higher in nitrogen than that of the native cells (detailed calculations provided in **Supporting Information**). The hydrogen produced in argon was higher than that in nitrogen, because nitrogenase can be regarded as a bifunctional enzyme under physiological conditions, where all the electrons delivered to the enzyme are used to release hydrogen in the absence of N_2_^.28^ Moreover, nitrogenase activity (measured by ethylene yield) in biohybrid system was higher than that in the native cells under both the N_2_ and Ar atmosphere (Fig. 2b). As previously reported, intracellular ammonium ions inhibited the nitrogenase activity,^29^ which explained the observations that the nitrogenase activity was higher in argon than in nitrogen. To further clarify the contributions of CdS-mediated photocatalytic water splitting to the overall hydrogen yield, we measured the hydrogen production of chemically synthesized CdS nanomaterial alone and found the hydrogen yield was very low (Fig. S4a). Moreover, the malic acid concentration in cell-free medium under photosynthetic conditions identical to those applied on biohybrid system decreased gradually at the presence of chemically synthesized CdS nanomaterial (Fig. S4b). These results show that the hydrogen produced by *R. rubrum*- CdS biohybrid system under photosynthetic conditions was predominantly contributed by the biohybrid *R. rubrum* cells, and the malic acid supplemented in culture medium served dual-purposes as both a nutrient substrate supporting cellular metabolism and a sacrificial agent being consumed under photosynthetic hydrogen evolution.

**Figure 2.**
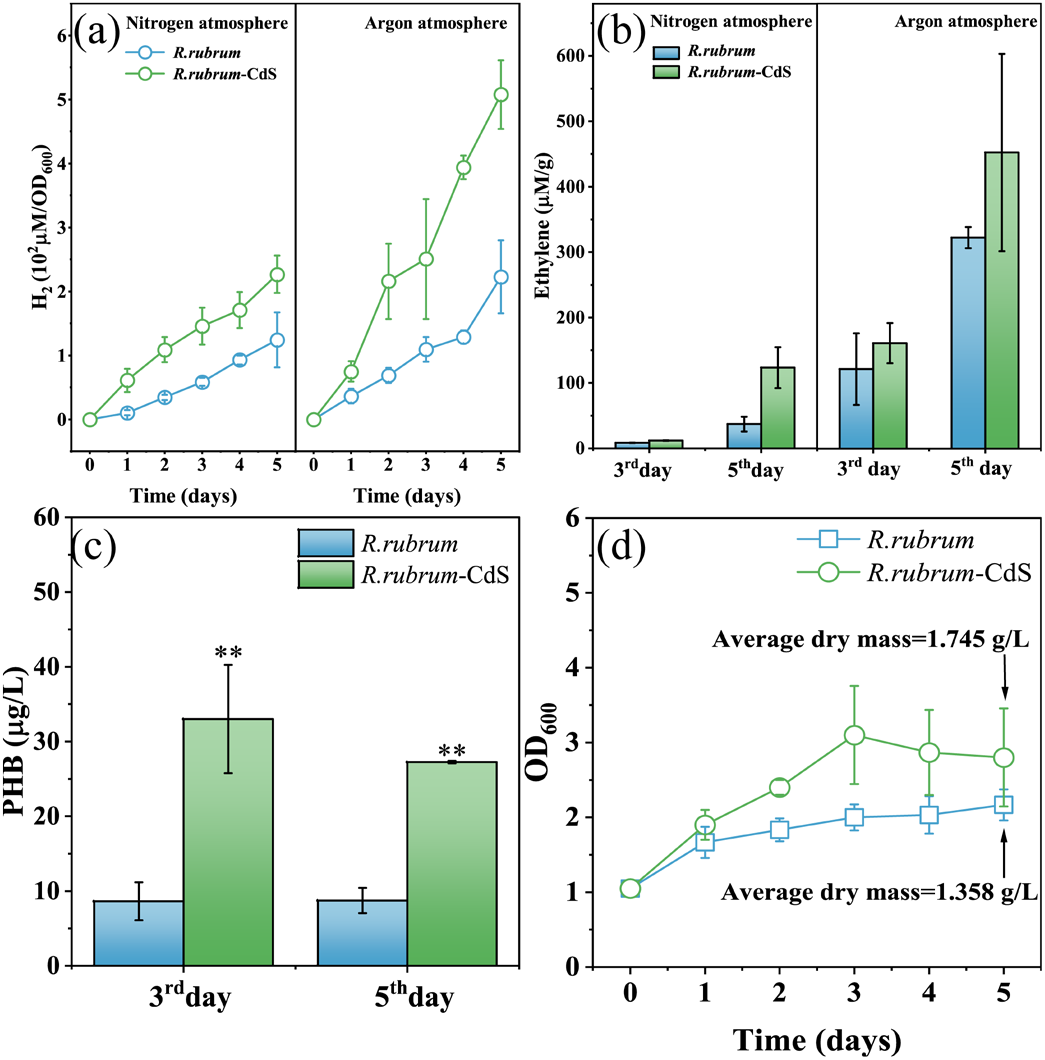
Metabolic profiles of R. rubrum-CdS biohybrid system: (a) H2 evolution in the headspace under N2 and Ar atmosphere. (b) Activity of nitrogenase in the headspace under N2 and Ar atmosphere. (c) PHB produced by native R. rubrum and R. rubrum-CdS biohybrid system at the 3^rd^ and the 5^th^ day. (d) Cell densities in the form of OD600 values and the final yield of average dry mass for cell pellets of the native *R. rubrum* and *R. rubrum*-CdS biohybrid system.

The intracellular PHB content (Fig. 2c) in biohybrid system (33.01 µg L^-1^) was 3.83-folds higher than that in the native cells (8.63 µg L^-1^) after 3 days’ culture, and such difference was reduced (27.26 µg L^-1^ for biohybrid system versus 8.73 µg L^-1^ for native cells) after 5 days’ culture, and the difference was statistically significantly (unpaired T-test, **p < 0.01). The growth profiles of native cells and biohybrid system were monitored via measuring the OD_600_ of both cultures at an interval of 24 h (Fig. 2d). In a 5-day period of continued culture, the biohybrid system underwent two distinct growth phases: a rapid growth from day 0 to day 3 (from OD_600_ = 1.0 at day 0 to OD_600_ = 3.0 at day 3), and a stationary period during the remaining days (minor decrease of OD_600_ value from day 3 to day 5). While the native cells also underwent growth from day 0 to day 3, the net increment of OD_600_ (from 1.0 at day 0 to 2.0 at day 3) was smaller than that of the biohybrid system. The difference in growth rate from day 0 to day 3 also resulted in higher dry mass of cell pellet for biohybrid system (1.745 g L^-1^) than that of the native cells (1.358 g L^-1^) at the 5^th^ day of continued culture, and the difference was statistically significantly (unpaired T-test, **p < 0.01).

### 2.2 Mechanisms of light-augmented metabolic activities

We further studied the underlying mechanisms for light- augmented metabolic activities of *R. rubrum*-CdS biohybrid system. Firstly, the malic acid concentrations in culture media were measured (Fig. 3a). Consistent with the rapid growth of *R. rubrum* from day 0 to day 3 (logarithmic phase), the malic acid concentration rapidly declined, and the net consumption was lower for biohybrid system than for the native cells. During the period of day 3 to day 5 (stationary phase), the bacterial density did not increase, and the consumption of malic acid for both the native and biohybrid system slowed down and reached the levels similar to each other. The intracellular levels of photosynthetic pigments (Bchl a and carotenoids) and NADPH, the reducing equivalent directly generated from the photosynthetic light reactions, were analyzed, since they are key parameters to characterize the activities of solar-capture and subsequent light-to-chemical energy conversion.^30 31^ During the whole growth cycle, although the pigments content of *R. rubrum* and *R. rubrum*-CdS biohybrid system increased with time, there was no significant differences in pigment levels between them (Fig. 3b-c). No significant difference in intracellular NADPH levels can be identified between the native cells and the biohybrid system until day 3. However, biohybrid system had a substantially increased intracellular NADPH from day 3 to day 5, whereas the native cells had a rapid decrease in intracellular NADPH during the same period (Fig. 3d). In addition, levels of intracellular ATP and ammonium ion were measured on day 3 and day 5. On the third day, the ATP level of biohybrid system was higher than that of the native cells. On the fifth day, the intracellular ATP level of the hybrid system decreased (Fig. 3e). The ammonium ion concentration of the biohybrid system was higher than that of the native cell on the third and the fifth day, and the total ammonium ion concentration of both the native and biohybrid cells decreased on the fifth day. Similarly, more rapid consumption of ammonium ion than nitrogen have been discovered on nitrogen-fixing bacteria in earlier studies^32^(Fig. 3f). We attribute these results to the different ways in which the photogenerated electrons produced by CdS are utilized during the different growth phases of bacteria. In logarithmic phase, CdS-generated electrons are mainly used for rapid proliferation of bacterial cells and accumulation of metabolites. As an energy currency, the ATP of the biohybrid system was higher than that of native *R. rubrum*. During the stationary phase, metabolic activity of *R. rubrum* cells slowed down for generating reducing power, a process that consumes ATP.^33^ The solar-capture function in biohybrid cells was carried out by CdS nanoparticles instead of the photosynthetic pigments, thus they were less likely to further modulate their intracellular levels and gave rise to the observations presented in Fig. 3b, and 3c.

**Figure 3.**
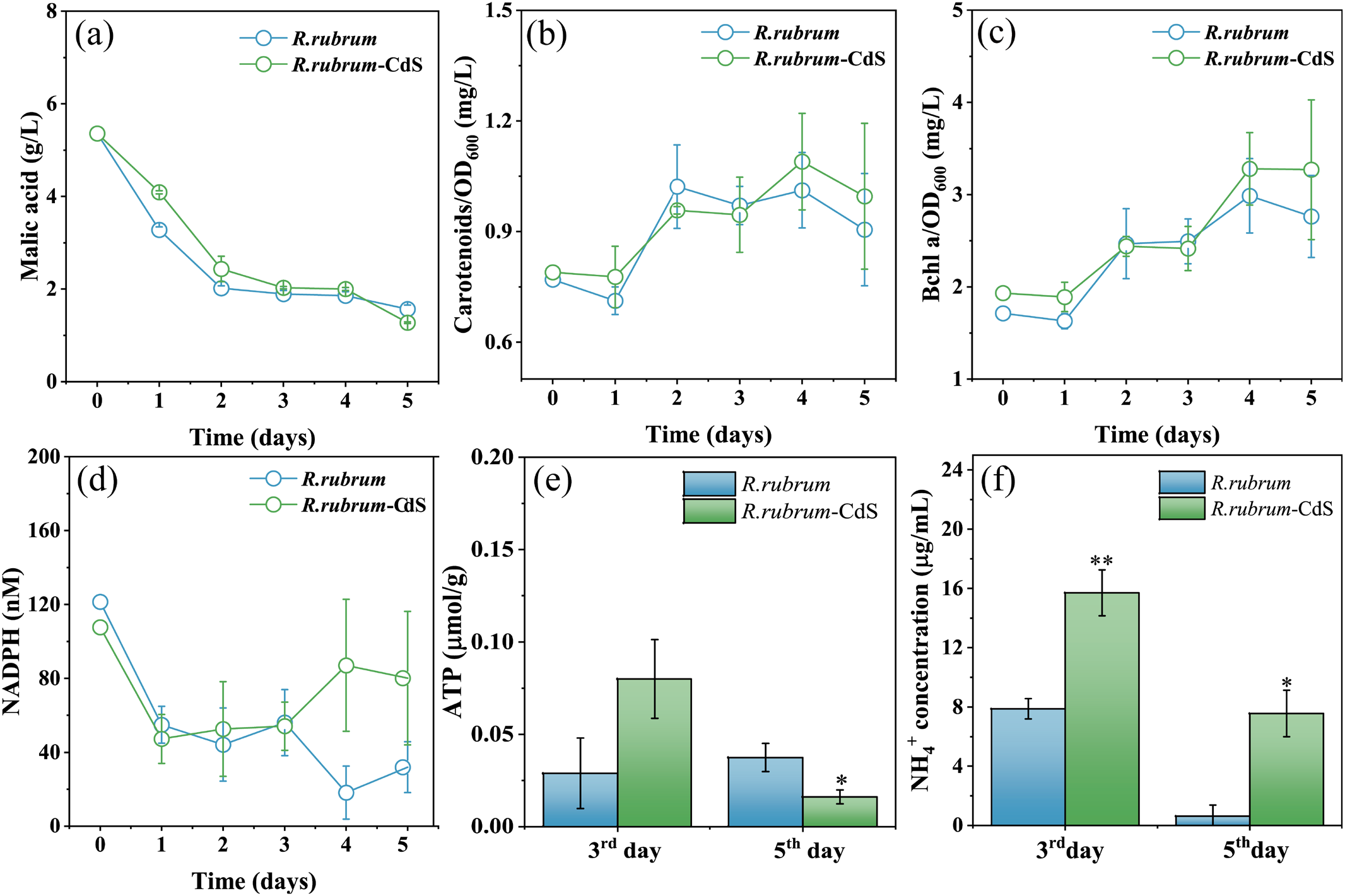
Light-augmented metabolic activities. (a) malic acid concentration in MMG medium. (b) intracellular Carotenoids content. (c) intracellular Bacteriochlorophyll a (Bchl a) content. (d) intracellular NADPH content. (e) intracellular ATP content.

### 2.4 Electron transfer mechanism between nanomaterials and purple bacteria

To understand the photoelectric conversion ability of *R. rubrum* and *R. rubrum*-CdS biohybrid system, we measured the cathode current and anode current of *R. rubrum* and *R. rubrum*-CdS biohybrid system, respectively. Within the potential range of 0 to - 1 V and starting from - 0.4 V, the difference in current density under light and dark conditions began to increase. Starting from -0.6V, the photocurrent of both systems gradually stabilized. With the increase of electric potential, the cathode current of *R. rubrum* was significantly higher than that of *R. rubrum*-CdS biohybrid system (Fig. 4a). The potential of - 0.6 V was selected as the initial voltage to measure the photocurrent densities of *R. rubrum* and *R. rubrum*-CdS biohybrid system, and found that *R. rubrum* gave rise to the photocurrent density significantly higher than that of the *rubrum*-CdS biohybrid system did (Fig. 4b), and *R. rubrum* exhibited a tendency of being the electron acceptor. Within the potential range of 0 to 1 V and starting from 0.4 V, the difference in current density under light and dark conditions began to increase, and the anode current of the *R. rubrum*-CdS biohybrid system was higher than that of *R. rubrum* (Fig. 4c). The potential of 0.6 V was selected as the initial voltage to measure the photocurrent density of *R. rubrum* and *R. rubrum*-CdS biohybrid system (Fig. 4d) and found the photocurrent density of *R. rubrum* was lower than that of *R. rubrum*-CdS biohybrid system, suggesting the *R. rubrum*-CdS biohybrid system behaved as an electron donor.

**Figure 4.**
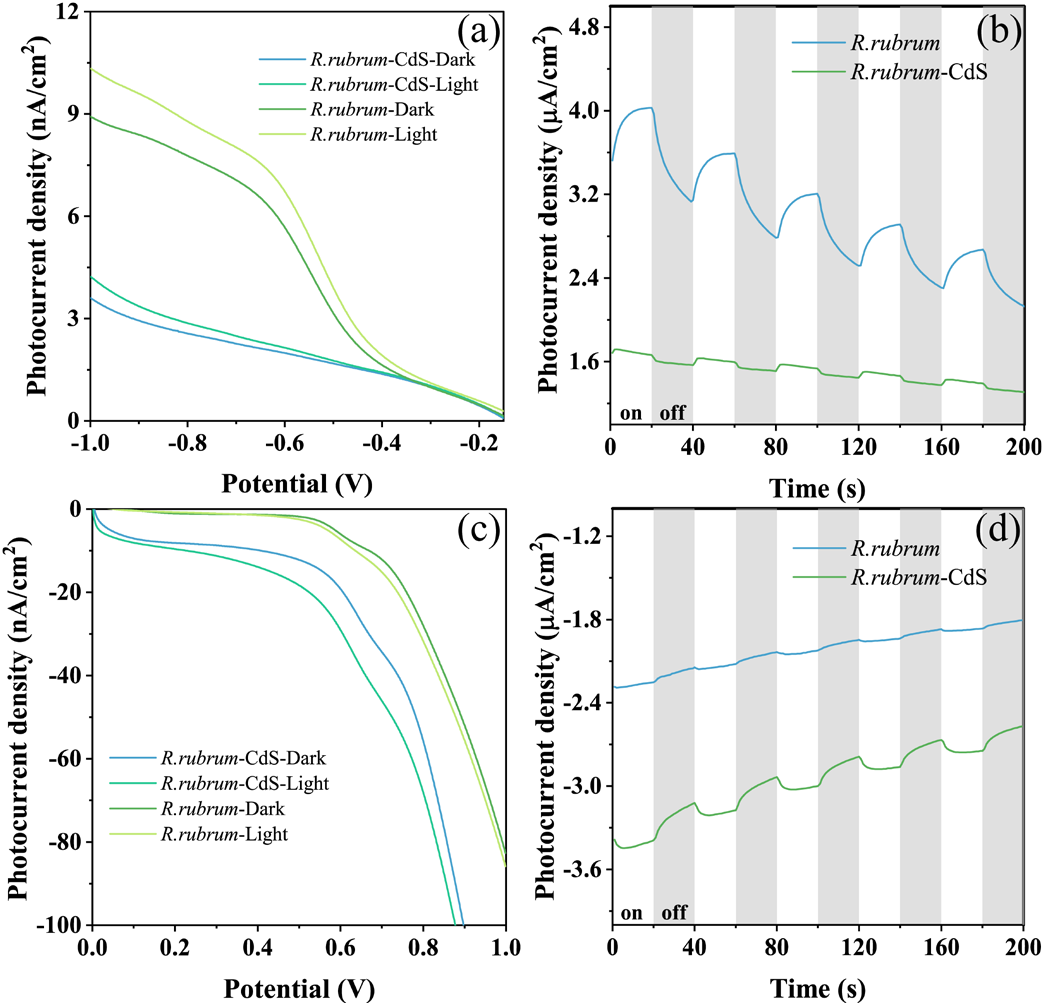
Photoelectrochemical analysis on *R. rubrum* and *R. rubrum*-CdS biohybrid system under white light. (a) LSV (0 to -1 V) curves of *R. rubrum* and *R. rubrum*-CdS biohybrid system. (b) photocurrent densities at -0.6 V. (c) LSV (0 to 1 V) curves of *R. rubrum* and *R. rubrum*-CdS biohybrid system. (d) photocurrent densities at 0.6 V.

To understand the light absorption and conversion capability of *R. rubrum*-CdS biohybrid system under monochromatic light, we chose blue light and red light to conduct photoelectrochemical studies on the biohybrid system. Starting from 0.4 V, the difference in current density between native R. rubrum cells and *R. rubrum*-CdS biohybrid system began to increase, which indicated that the electron-donating ability of *R. rubrum*-CdS biohybrid system under blue light was stronger than that under red light (Fig. 5a). We measured and compared the photocurrent densities of native *R. rubrum* cells and *R. rubrum*-CdS biohybrid system under blue light irradiation at the initial potential of 0.6 V, the results showed that the photocurrent densities of *R. rubrum*-CdS biohybrid system were higher than that of native *R. rubrum* cells (Fig. 5b). We also performed the LSV test under red light irradiation. At the potential of 0.8 V, the electron-donating ability of native *R. rubrum* cells was slightly higher than that of *R. rubrum*-CdS biohybrid system, and this trend was reversed above 0.8 V (Fig. 5c). We measured and compared the photocurrent densities of native *R. rubrum* cells and *R. rubrum*-CdS biohybrid system under red light at the initial potential of 0.6 V. the results showed that the photocurrent level had little difference (Fig. 5d).

**Figure 5.**
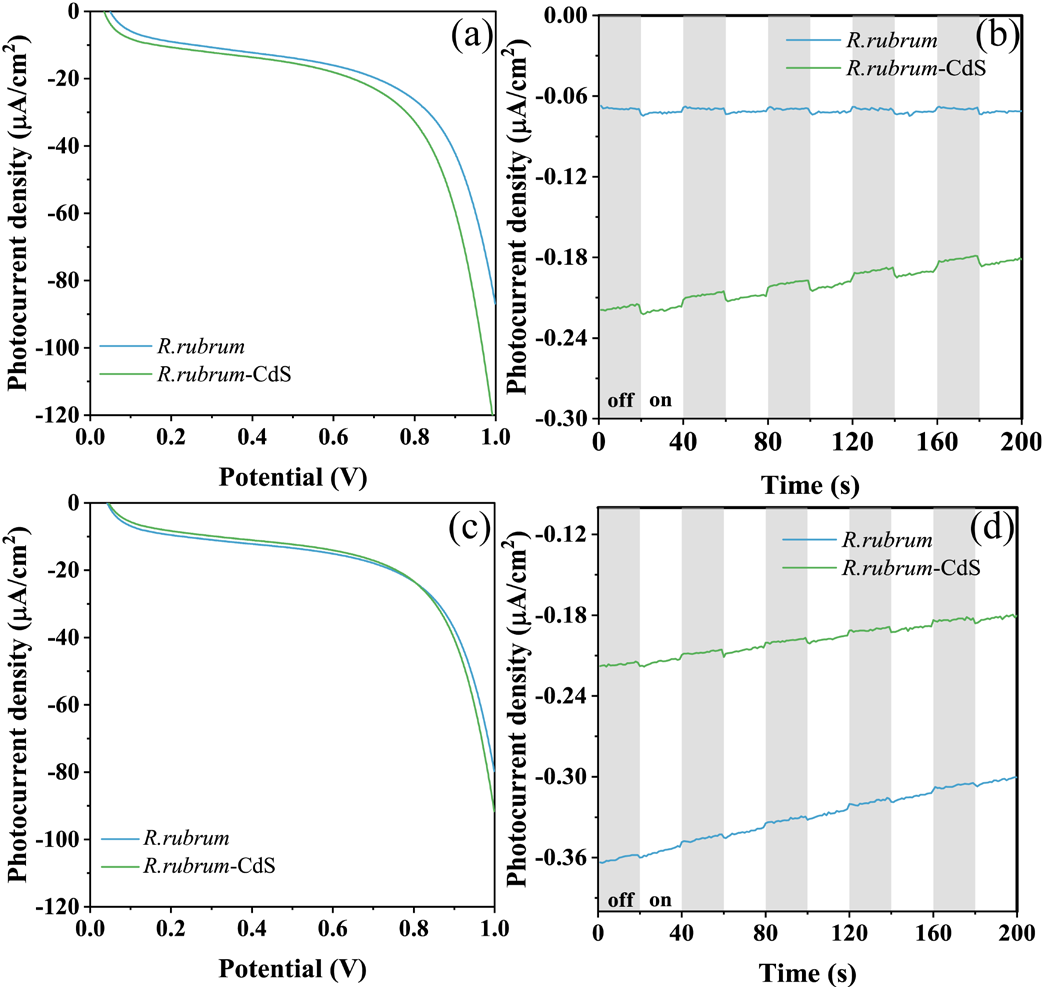
Photoelectrochemical analysis on *R. rubrum* and *R. rubrum*-CdS biohybrid system under monochromatic light. (a) LSV (0 to 1 V) curves of native *R. rubrum* cells and *R. rubrum*-CdS biohybrid system under blue light irradiation. (b) photocurrent densities of native *R. rubrum* cells and *R. rubrum*-CdS biohybrid system under blue light irradiation. (c) LSV (0 to 1 V) curve of native *R. rubrum* cells and *R. rubrum*-CdS biohybrid system under red light irradiation. (d) photocurrent densities of native *R. rubrum* cells and *R. rubrum*-CdS biohybrid system under red light irradiation.

To understand the electron transfer mechanism at the material- cell interface, we tested time-resolved PL spectra and performed electrochemical impedance analysis on the biohybrid system. The average lifetime of the photoelectrons in the *R. rubrum*-CdS biohybrid system was estimated to be about 643 ps, which was longer than the chemically synthesized CdS (631 ps) (Fig. 6a). These results suggested that the photogenerated electron−hole pairs of biomineralized CdS nanoparticles were efficiently separated, and an electron transfer pathway between surface-deposited CdS and *R. rubrum* was established in the *R. rubrum*-CdS biohybrid system.^34^ The electrochemical impedance spectra of native *R. rubrum* and *R. rubrum*-CdS biohybrid system were fitted with ZSimpWin software (fitting error 10^−3^ and 10^−4^), the equivalent circuit diagram is also presented (Fig. 6b, parameters provided in figure caption). The good stability of this system was demonstrated by the stable Rs value. The low-frequency region impedance (Re) corresponded to the impedance within the biofilm.^35^ CdS nanoparticles on cell membrane formed a convex surface to reduce the distance of electron transfer within the biofilm. Under the irradiation of white light, the low-frequency impedance spectra of biohybrid system had smaller radius than those of the native cells, which was opposite to those observed under dark (Fig. 6c). This observation confirmed that light-excited charge separation of CdS enhanced electron flow between cells and electrode. The intermediate frequency region corresponded to the Warburg impedance. The high-frequency region of the impedance spectra was fitted with semicircles, whose radius were proportional to the conduction resistance (Rct). Electron transduction corresponding to the high-frequency region usually occurs between the electrode and the double layer of solution, which were not different significantly from each other under different conditions (Fig. 6d). The low-frequency region impedance (Re) was lower under the blue light than under the red light, which was also due to the generation of photoelectrons under blue light irradiation (Fig. 6e). The results under the red light were potentially due to its lower energy (approximately 2.0 eV for 620 nm) than the bandgap of CdS (2.37 eV), thus the red light was unable to excite the charge separation of CdS. Therefore, unexcited CdS nanoparticles affected the current flow between *R. rubrum* cells and electrode, which resulted in the higher impendence for biohybrid system than for native cells. Similar to those observed under white light irradiation, there were no significant differences for the electron transduction between the electrodeand the double layer of solution within the high-frequency region under the irradiation of red and blue light (Fig. 6f).

**Figure 6.**
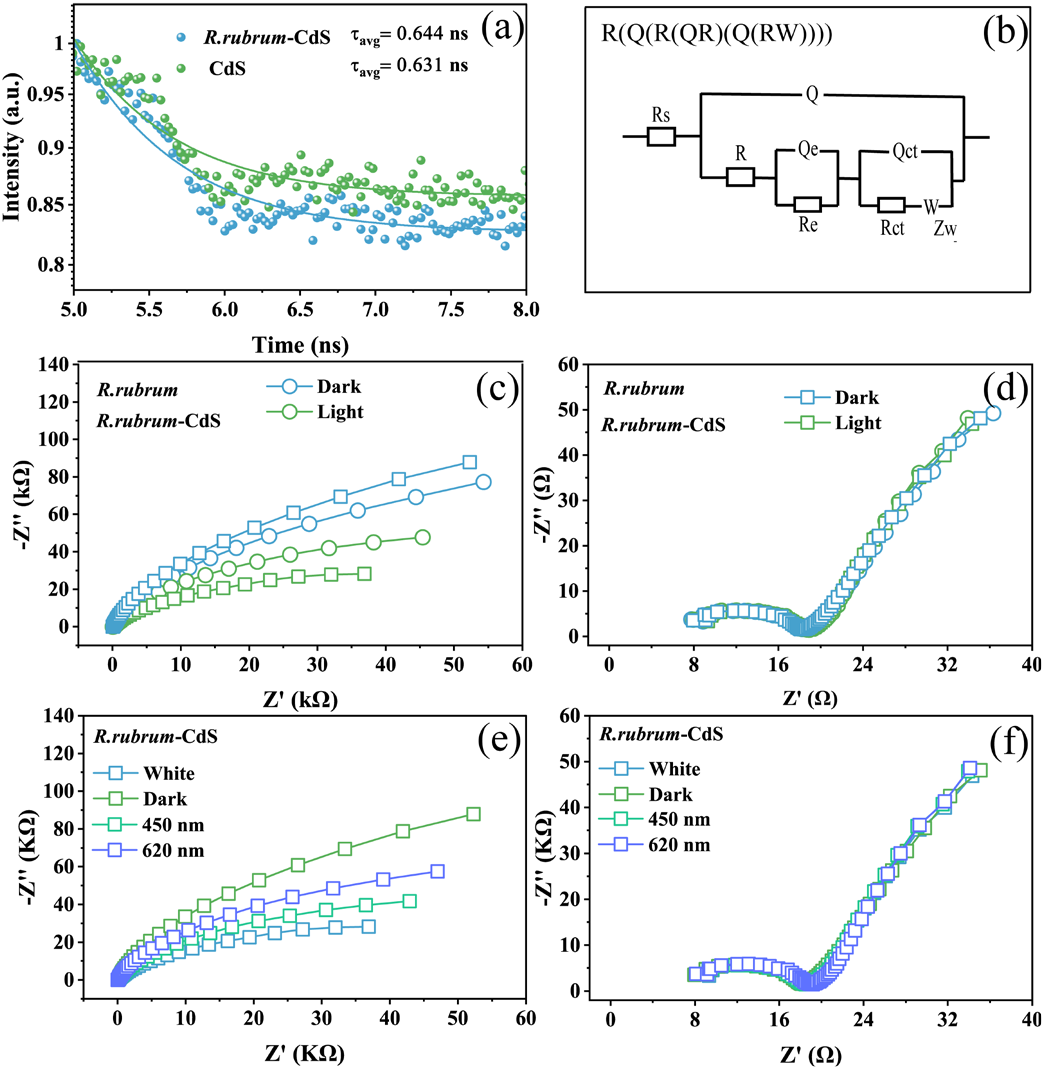
Electron transfer ability of interface: (a) Time-resolved PL spectra of *R. rubrum*- CdS and CdS compound (b) Equivalent circuit diagram of native *R. rubrum* cells and *R. rubrum*-CdS biohybrid system. Parameters for equivalent circuit diagram: Rs, solution resistance; R, bacterial resistance; Rct, conduction resistance; Qct, capacitance of electric double layer; W, Warburg impedance; Re, low-frequency region impedance; Qe, capacitance of biolilm; Q, total capacitance (c) fitted electrochemical impedance spectra of native *R. rubrum* cells and *R. rubrum*-CdS biohybrid system under dark or white light irradiation. (d) zoom-in view of the high-frequency region of Fig 6c. (e) fitted electrochemical impedance spectra of *R. rubrum*-CdS biohybrid system under the irradiation of white, blue, and red light. (f) zoom-in view of the high-frequency region of Fig 6e.

## 3. Conclusions

Compared to the native cells under identical culture conditions, biohybrid system gave rise to increased H_2_ evolution and nitrogenase activity, higher intracellular PHB content, and more rapid cellular growth and biomass accumulation. These findings proved that the CdS nanoparticles augmented the general cellular metabolic activity of *R. rubrum* cells.

We have analyzed the time-dependent changing in photosynthetic (Bchl a and carotenoids), energetic (intracellular NADPH, ATP), and metabolic (malic acid consumption and intracellular NH_4_^+^) parameters of native cells and biohybrid system, and discovered that these parameters also followed the “two-phases” pattern that have been identified from the growth experiment. These results underlined that different bacterial growth phases determined the modes of energy utilization and metabolic profiles of *R. rubrum* cells. The *R. rubrum* cells under photoheterotrophic growth acquired energy from both the organic substrate (i.e., malic acid) and the solar power.^36^ In the case of *R. rubrum*-CdS biohybrid system, the solar capture was synergistically carried out by both the native photosystem and the CdS nanoparticles, the increased energy supply to the bacterial metabolic system thus augmented the activity of nitrogenase and the growth rate from day 0 to day 3. The faster energy-to-biomass turnover of biohybrid system also resulted in higher intracellular ATP and PHB content. These results suggested CdS-generated photoelectrons provide additional energy for the growth of *R. rubrum*.

**Figure 1a.**
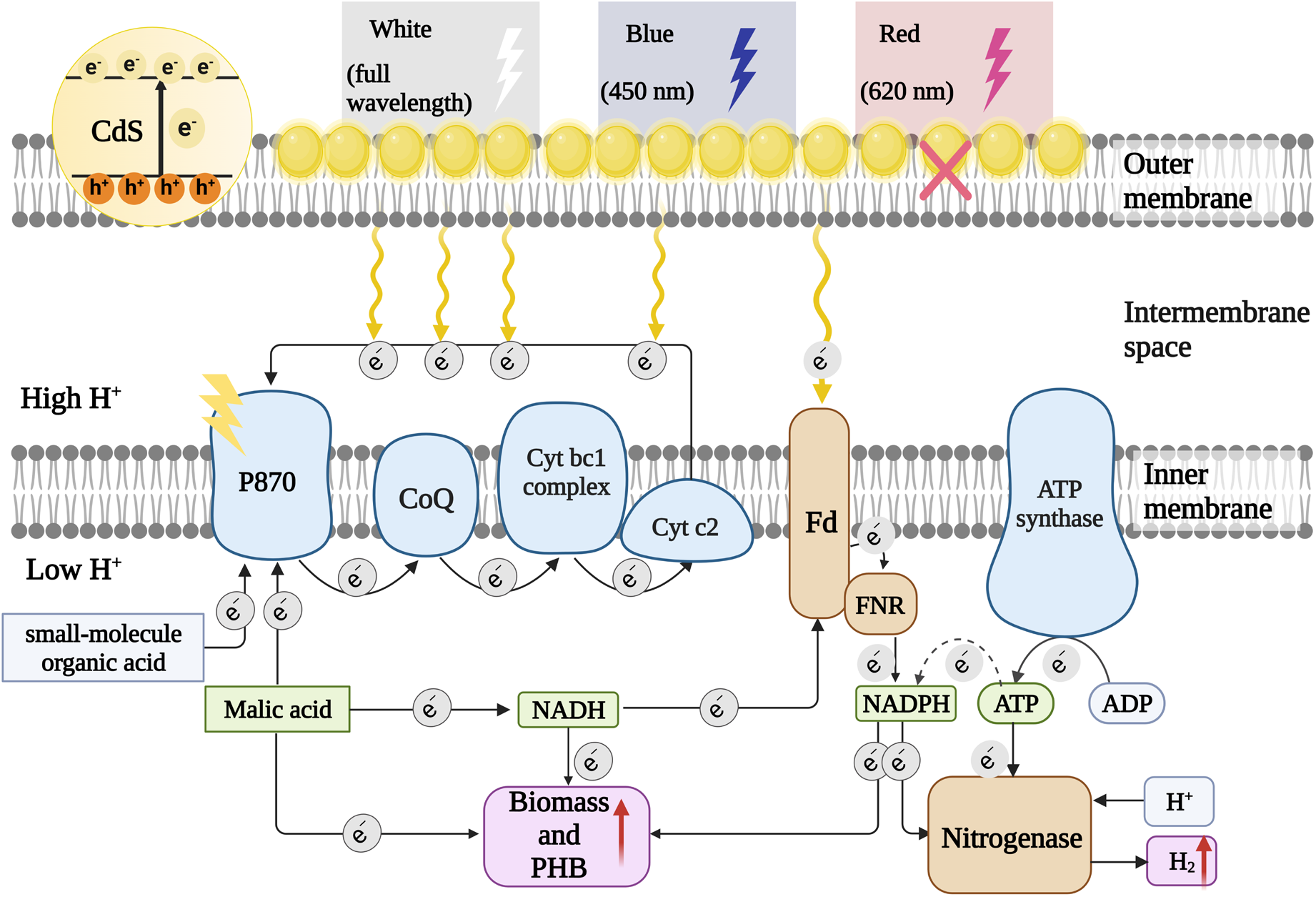
Proposed electron flow of augmented metabolism for *R. rubrum*-CdS biohybrid system. Abbreviations: P870, pigment 870; CoQ, coenzyme Q; cyt bc1 complex, cytochrome bc1 complex; cyt c2, cytochrome c2; Fd, Ferredoxin; FNR, Ferredoxin-NADP+ Reductase

During the stationary phase, the effect of CdS-augmented energy metabolism was underlined by a number of cellular energetic parameters other than the rapid growth, i.e., intracellular NADPH levels. The increased levels of NADPH implied that biohybrid *R. rubrum* cells up-regulated the photosynthetic activity to enhance solar energy utilization, an action that effectively countered the reduced supply of organic energy source (below 2 g/L malic acid from day 3 to day 5). Overall, the energy metabolic profile of native cells and biohybrid system were consistent with the pattern of growth phases, and the CdS-augmented cellular metabolism was underlined by the disparity between the native cells and biohybrid system on various metabolic parameters of metabolites.

Based on the results presented above, we outlined the mechanism underlying the augmented photosynthetic activities of *R. rubrum*-CdS biohybrid system^37^ (Fig. 7). The surface-deposited CdS nanoparticles generated photoelectrons under visible light irradiation, which entered the P870 electron conduction chain and ferredoxin (Fd).^38 39^ The Fd delivered electrons to nitrogenase, and part of them were utilized to boost H_2_ evolution, which also verified the increased nitrogenase activity and the subsequently increased biomass accumulation. Another part of photoelectrons was used to power the regeneration of intracellular NADPH and ATP, which demonstrated that the R. rubrum cells with surface-deposited CdS nanoparticles were more productive in H_2_ generation, PHB production and biomass accumulation than those of the native cells.

In conclusion, we successfully constructed *R. rubrum*-CdS biohybrid system via a biomineralization strategy and evaluated its photosynthetic activity. The elevated yields of metabolic products confirmed the augmented metabolic activities of biohybrid system over its native counterpart that resulted from the CdS-supplied photoelectrons. We also conducted electrochemical and monochromatic light excitation experiments on the biohybrid system to delineate its light absorption and conversion profiles and the electron transfer properties at material-cell interface. The results demonstrated that the formation of *R. rubrum*-CdS biohybrid system fostered the light utilization of the entire system, which further confirmed the promising prospects for utilizing non-oxygenic photoheterotrophic microbials in the field of semi-artificial photosynthesis. The biohybrid cells gave rise to growth-phase dependent metabolic and energy consumption profiles that were different from those of the native cells, which highlighted the CdS nanoparticle could modulate the metabolism of the host cells via supplying additional photoelectrons. In addition, the use of electrochemical techniques to investigate the electron transfer properties at the material-bacteria interface provided a good reference for understanding the interactions between purple bacteria and nanomaterials in biohybrid systems.

## 4. Experimental

### 4.1 Construction of *R. rubrum*-CdS biohybrid system

Biomineralization strategy was applied to obtain the biohybrid system of *R. rubrum-*CdS. The stock suspension of *R. rubrum* (ATCC11170) was centrifuged (6000 g, 5 min), and the bacterial cells were inoculated in MMN medium (Table S1) for activation, and the initial cell density (OD_600_) was adjusted to 1. The cells were cultured to the mid-log phase, centrifuged (6000 g, 5 min) and washed three times with phosphate buffered saline (PBS, pH = 7), resuspended in MMN medium to OD_600_ = 1, and aliquoted into 250 mL anaerobic bottles. The aliquots were supplemented with *L*-cysteine hydrochloride (1 mM) and different concentrations of Cd(NO3)2 (0.05 mM, 0.1 mM, 0.2 mM, 0.3 mM, respectively), mixed well and cultured in a water bath under the illumination of LED (temperature at 30 °C, light intensity at 30 mW/m2, illumination area 0.002 m2, magnetic stirring at 200 rpm). After 24 h of incubation, the samples for scanning electron microscopy (SEM) analysis were prepared as the following method: bacterial suspension (1 mL) was centrifuged (13000 g, 5 min) and washed three times with PBS, and fixed with glutaraldehyde (4%, 1 mL) at 4 °C for 2 h, and then washed twice with PBS. The fixed samples were then dehydrated sequentially with ethanol/water mixtures of gradient concentrations (50%, 75%, 85%, 95% and 100%), dripped onto the polished surface of silicon wafer and air-dried. Samples were analyzed with a desktop field-emission SEM (Phenom Pharos-SED-EDS G2, Thermo Fisher) at an acceleration voltage of 15 kV, the elemental distribution maps of the samples were analyzed with energy-dispersive X-ray spectroscopy (EDS) using an Amptek Fast SDD X123 Ultra High-Performance Silicon Drift Detector at an excitation voltage of 15 kV.

### 4.2 Photosynthetic activities of biohybrid system

The *R. rubrum* suspension was cultured for 24 h, centrifuge (6000 g, 5 min) and washed three times with PBS, and diluted to OD_600_ = 1 with MMG medium (Table S2). Aliquots of the suspensions (100 mL) were filled into the 250 ml anaerobic bottles, the headspace was purged with nitrogen, and the bottle were sealed tight. The suspension was incubated for 5 days, samples were taken every 24 h and analyzed.

### 4.3 Hydrogen evolution

A microinjection needle was used to take gas samples (100 μL, each) from the headspace and manually injected into a gas chromatograph (GC9790II., Fuli, China) to analyze the levels of hydrogen in samples. Quantification of hydrogen was based on the external standard method, the carrier gas was argon (purity > 99.99%), the flow rate was 30 mL min^-1^, the detector was a thermal conductivity detector (TCD), the column chamber temperature was 80 °C, and the retention time was 5 min.

### 4.4 Cell density and dry mass

The OD_600_ values of *R. rubrum* suspension was determined every 24 h using a cell density meter (WPA CO8000, Biochrom, UK). For the measurement of dry mass, the bacterial suspensions (50 mL each) at the beginning of the experiment and at the 5^th^ day of culture were collected and centrifuged (13000 g, 10 min). The supernatant was discarded, the cell pellets were snap-frozen in liquid nitrogen and lyophilized for 48 h (Scientz-10N/A, SCIENTZ, China), and their mass were measured.

### 4.5 Nitrogenase activity

The bacterial suspensions in reaction bottles were purged with Ar or N_2_ to remove the residual H_2_. A 10% volume of the headspace gas was replaced with equal volume of acetylene (C_2_H_2_). The reaction bottles were sealed and placed into a light incubator for 1 h before the analysis of ethylene yield. A microinjection needle was used to take gas samples (100 μL, each) from the headspace and manually injected into a gas chromatograph (GC9790II., Fuli, China) to analyze the levels of ethylene in samples. Quantification of ethylene was based on the external standard method, the carrier gas was argon (purity > 99.99%), the flow rate was 30 mL min^-1^, the detector was a flame ionization detector (FID), the column chamber temperature was 80 °C, and the retention time was 20 min.

### 4.6 PHB quantification

The lyophilized *R. rubrum* pellets were digested with concentrated sulfuric acid (98%, 1 mL) in glass tubes and heated in water bath (90 °C, 30 min). The digested mixture was diluted with ultrapure water (4 mL) and filtered through 0.22 μm membrane. The PHB concentration in these samples were measured using a high performance liquid chromatography (1260 Infinity II, Agilent, USA) under the following parameters: the mobile phase was 5 mM sulfuric acid, the column was HPX- 87H (Aminex, Bio-rad, USA), the flow rate was 0.6 mL min^-1^, the injection volume was 10 μL, the column temperature was 60 °C, the diode array detector (DAD) was set to the wavelength at 210 nm.

### 4.7 Malate quantification

The reaction samples (1 mL) were taken every 24 h. They were centrifuged (13000 g, 5 min), the supernatant was collected and filter through 0.22 μm membrane. The malate concentration in these samples were measured using the same high performance liquid chromatography and column as those for the PHB quantification. The refractive index detector (RID) was set to 55 °C, other parameters were identical to those for PHB quantification, and the retention time was 9 min.

### 4.8 NADPH quantification

Quantification of NADPH in reaction samples was performed using the NADP^+^/NADPH Assay Kit (WST-8 method, Cat. No. S1079, Biyuntian, China). The sample suspensions (200 μL) from day 1 to day 5 were collected and centrifuged (12000 g, 5 min). The supernatant was discarded, and the cell pellets were lysed in extraction buffer (200 μL). The samples were centrifuged (12000 g, 4 ºC, 5 min), the supernatant was collected (100 μL) and heated at 60ºC for 30 min, and centrifuged again (12000 g, 4 ºC for 5 min), and the supernatant (50 μL) was added into the wells of a 96-well plate. The G6PDH working solution was added (100 μL per well), and the plate was incubated under dark at 37ºC for 10 min. The color development solution was added (10 μL per well), and the plate was incubated at 37 ºC for 10-20 min to facilitate the formation of an orange-yellow formazan solution, and the absorbance at 450 nm was measured using a microplate reader (SpectraMax iD3, Molecular Devices, Austria). The NADPH concentrations in samples were calculated according to the standard curve.

### 4.9 Photosynthetic pigments quantification

The photosynthetic pigments were extracted using an extraction mixture of methanol:acetone = 2:7 (v/v). The reaction suspension (1 mL) was collected and centrifuged (6000 g, 5 min). The supernatant was discarded, and the cell pellets were washed twice with ultrapure water. The cell pellets were extracted repeatedly with extraction mixture (1 mL) until the they turned white. The samples were centrifuged (13000 g, 5 min), the supernatant was added to the wells of a 96-well plate, and the absorbance at 496 nm, 515 nm, and 763 nm were measured with a microplate reader (SpectraMax iD3, Molecular Devices, Austria). The pigment concentrations were calculated according to the following formula^40^(Bchl a, Bacteriochlorophyll a):

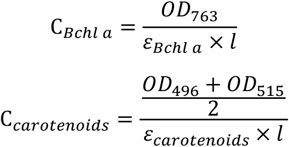

Where ε _Bchl a_=82.2 g L^-1^ cm^-1^, ε _carotenoids_ = 250 g L^-1^ cm^-1^.

### 4.10 ATP quantification

Quantification of ATP in reaction samples was performed using the ATP Assay Kit (Cat No. S0026, Biyuntian, China). The sample suspensions (200 μL) of day 3 and day 5 were collected and centrifuged (12000 g, 5 min). The supernatant was discarded, and the cell pellets were lysed in extraction buffer (200 μL). The supernatant was collected (100 μL) and mixed with detection solution (100 μL), and the luminescence was measured using a microplate reader (SpectraMax iD3, Molecular Devices, Austria). The ATP concentrations in samples were calculated according to the standard curve.

### 4.11 NH_4_ ^+^ quantification

The sample suspensions (200 μL) of day 3 and day 5 were collected and centrifuged (12000 g, 5 min). The supernatant was discarded, and the cell pellets were washed in PBS twice. The supernatant was discarded and the cell pellets were lysed in lysis buffer (10% sucrose, 300 mM NaCl, 90 mM EDTA, 3-mg/mL lysozyme, 50 mM Tris-HCL [pH 7.5]).^41^ Next, the suspension was mixed and incubated on ice for 2 h. The cell suspension was then frozen and thawed five times by cycling between 37°C waterbath and liquid nitrogen. Finally, the suspension was sonicated (30% power) for 3 min using an Ultrasonic broken instrument. The sample suspensions were centrifuged (12000 g, 5 min), and the supernatant (20 μL) was mixed with solutionⅠ and solutionⅡ, left to stand at 25 °C for 1 hour, and mixed with solution Ⅲ. The absorbance at 625 nm were measured with a microplate reader (SpectraMax iD3, Molecular Devices, Austria).

### 4.12 Electrochemical studies

The *R. rubrum* suspension (40 μL of OD_600_ = 5) was uniformly applied on one side of the carbon paper (1 cm × 1 cm). The suspension was dried, and 20 μL of Nafion (0.05% in absolute ethanol) was added onto the carbon paper, which was then clamped with electrode clips. The working electrode (carbon paper with *R. rubrum* cells), reference electrode (Ag/AgCl reference electrode), and the counter electrode (Pt sheets) was installed into a quartz electrolyte cell (50 mL volume). The cell was filled with electrolyte solution (PBS, 40 mL), purged with nitrogen gas (10 min) and sealed. The light reactions were irradiated under the LED source of white light, or monochromatic light at 450 nm or 620 nm with bandpass filter, respectively (perfect light, China), the light intensity was 20k lux. The following photoelectrochemical analysis were performed on a electrochemical workstation (CHI760E, Shanghai Chenhua, China): (1) For linear-sweep voltammetry (LSV), the voltage range was 0 to -1 V and 0 to 1 V, the sweep speed was 0.01 V s^- 1^; (2) For amperometric-time (I-t) response, the current values at the -0.6 V and 0.6V potential (versus Ag/AgCl) were recorded; (3) For alternate-current AC impedance test, the initial voltage was the open circuit potential, the high frequency parameter was set to 10^6^ Hz, the low frequency parameter was set to 10^−2^ Hz, the AC impedance spectrum under this parameter was recorded, and the equivalent circuit diagram was fits using ZSimp Win software.

## Author Contributions

BW and CZ designed the project. LW, and SS, performed the experiments and collected the data. LW, CZ and JL analyzed and interpreted the data. BW, XX, CZ and JL edited, compiled, and finalized the draft. All authors contributed to the article and approved it for publication. #LW and SS contributed equally to this paper.

## Acknowledgements

This work was supported by the National Key R&D Program of China (Grant No. 2021YFA0910800), National Natural Science Foundation of China (Grant No. 42002303 and 22008252), Outstanding Youth Project of Guangdong Natural Science Foundation (Grant No.2022B1515020047), General project of Natural Science Foundation of Guangdong Province (Grant No.2021A1515010253), General project of Natural Science Foundation of Guangzhou (Grant No.202102020523). The authors thank to Prof. Hang Xing from Hunan University for providing the *R. rubrum* strain.

