## Supporting Information for "Semiconductor augumented valuable chemical photosynthesis from *Rhodospirillum rubrum* and mechanism study"

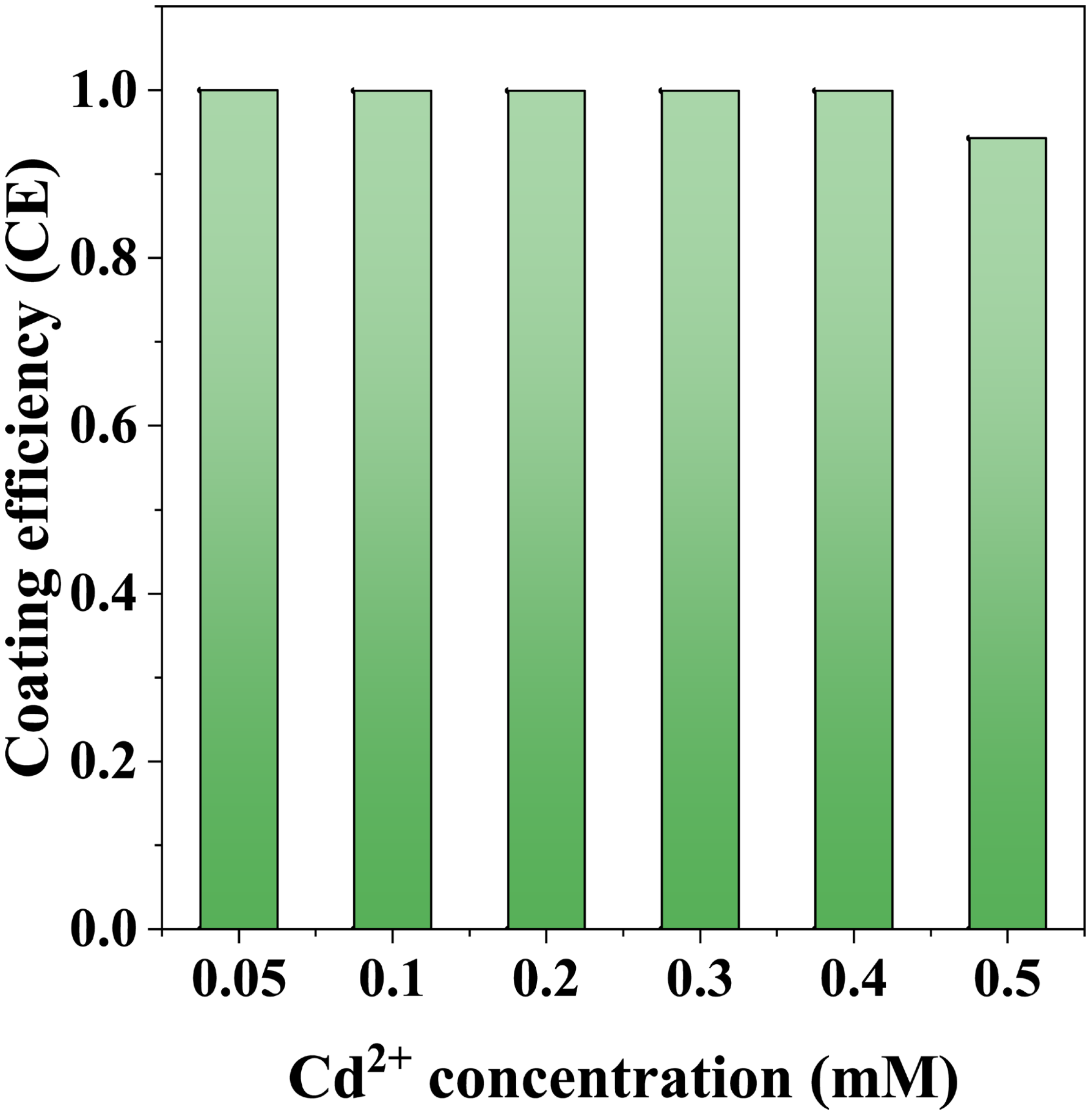


**Fig. S1** CdS coating efficiencies for biomineralization on *R. rubrum* cells using different concentrations of $Cd(NO_3)_2$ for 24 h

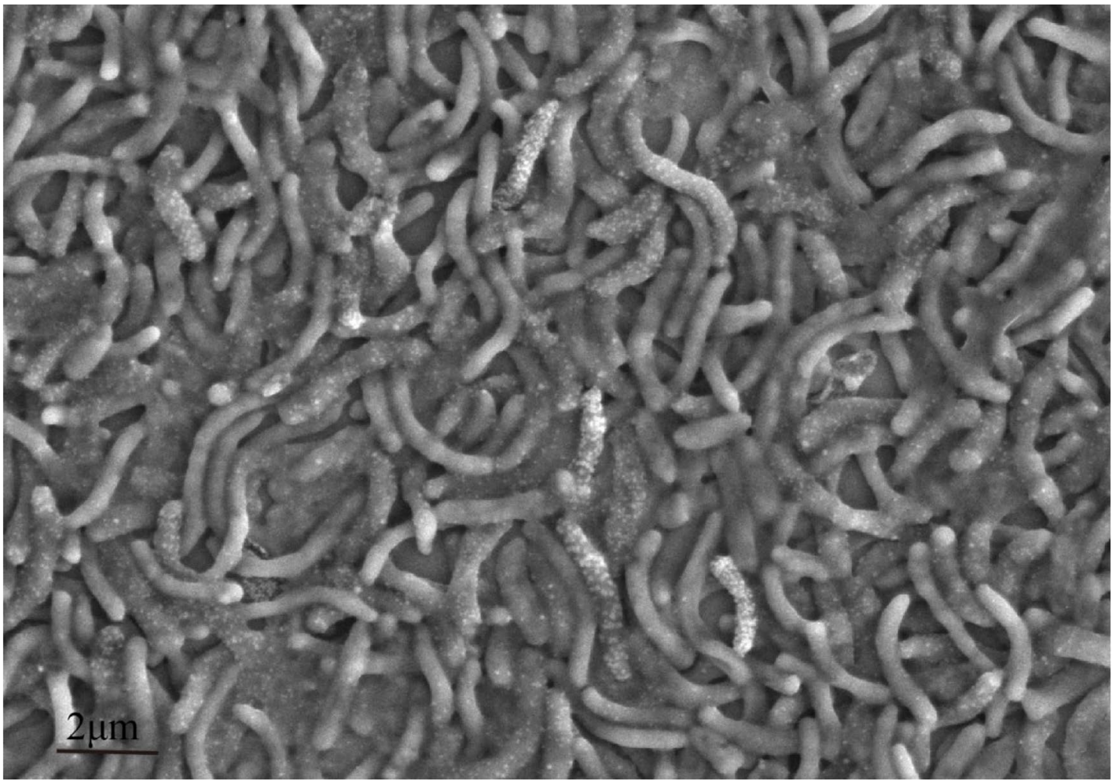


**Fig. S2** SEM image of *R. rubrum*-CdS biohybrid system prepared under a $Cd^{2+}$ concentration of 0.2 mM.

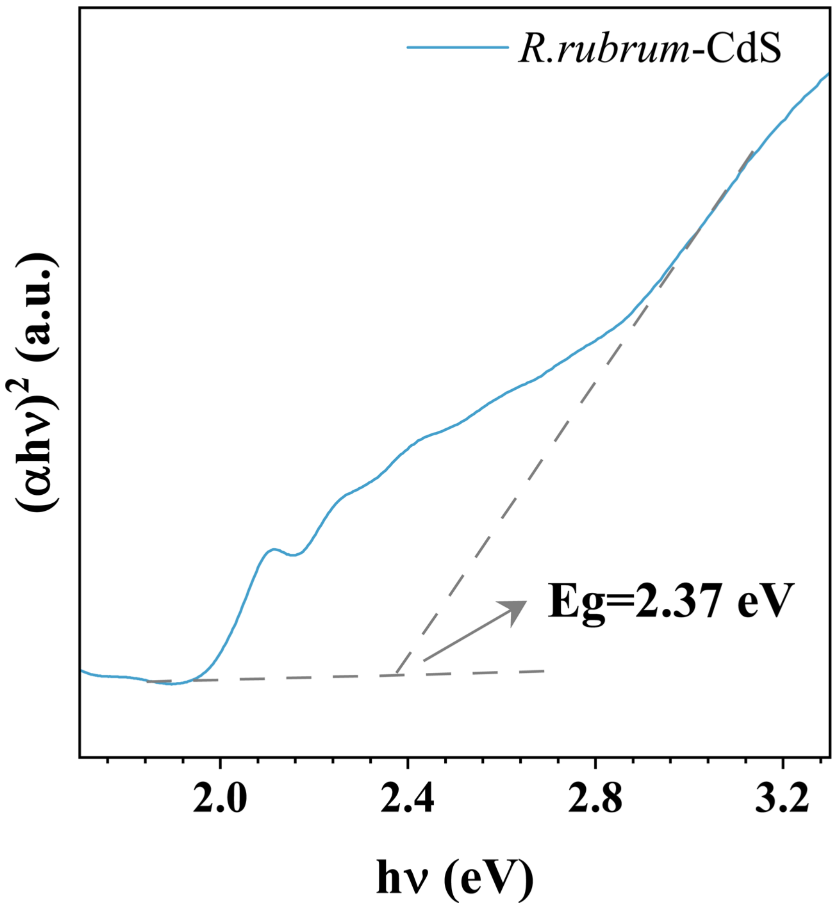


**Fig. S3** Tauc plots for band gap (*Eg*) determination of *R. rubrum*-CdS biohybrid system.

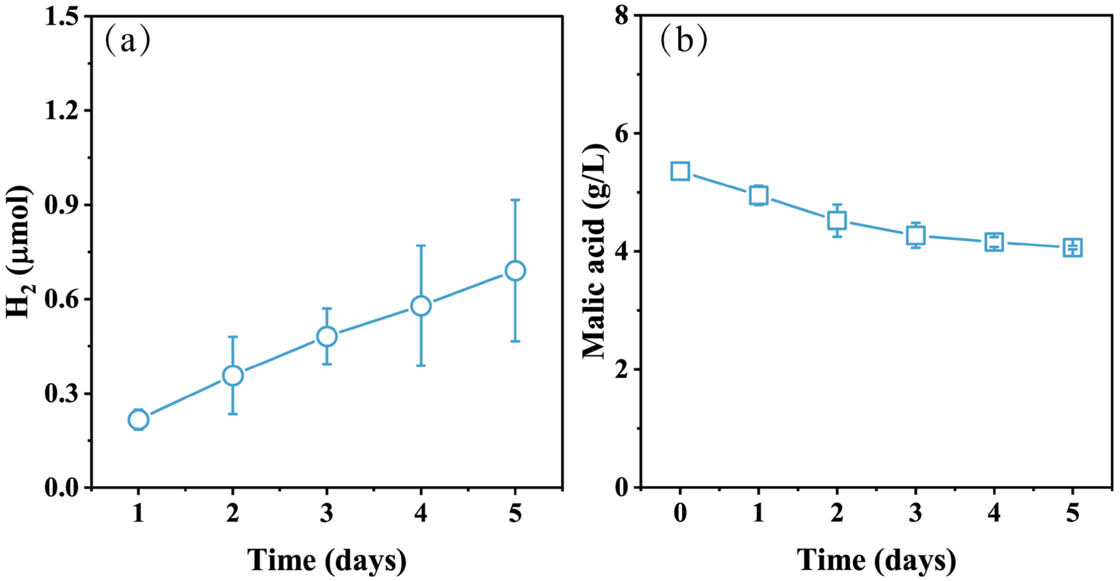


**Fig. S4 a** Photocatalytic $H_2$ evolution of chemically synthesized CdS. **b** Changes of malic acid concentration in cell-free MMG medium at the presence of chemically synthesized CdS.

**Table S1** Components of MMN medium

| MMN | 1000 mL |
|---|---|
| EDTA (2% w/v) | 1 mL |
| 4-Aminobenzoic acid (PAPA, 30 mM) | 0.1 mL |
| Malic acid | 5.36 g |
| $(NH_4)_2SO_4$ | 1.25 g |
| MOPS | 8.37 g |
| Tricine | 0.72 g |
| β-glycerophosphate·2Na | 0.344 g |
| $MgSO_4 \cdot 7H_2O$ | 0.2 g |
| $CaCl_2 \cdot 2H_2O$ | 75 mg |
| $FeSO_4 \cdot 7H_2O$ | 11.8 mg |
| $K_2SO_4$ | 20 mg |
| $H_3BO_3$ | 2.8 mg |
| $MnSO_4$·4H2O | 2.1 mg |
| $Na_2MoO_4 \cdot 2H_2O$ | 0.75 mg |
| $ZnSO_4 \cdot 7H_2O$ | 0.24 mg |
| $Cu(NO_3)_2 \cdot 3H_2O$ | 0.04 mg |
| KOH | Adjust pH to 6.8 |

**Table S2** Components of MMG medium

| **MMG** | **1000 mL** |
|---|---|
| EDTA (2% w/v) | 1 mL |
| 4-Aminobenzoic acid (PAPA, 30 mM) | 0.1 mL |
| Malic acid | 5.36 g |
| Glutamic acid | 2 g |
| MOPS | 8.37 g |
| Tricine | 0.72 g |
| β-glycerophosphate·2Na | 0.344 g |
| $MgSO_4 \cdot 7H_2O$ | 0.2 g |
| $CaCl_2 \cdot 2H_2O$ | 75 mg |
| $FeSO_4 \cdot 7H_2O$ | 11.8 mg |
| $K_2SO_4$ | 20 mg |
| $H_3BO_3$ | 2.8 mg |
| $MnSO_4 \cdot 4H2O$ | 2.1 mg |
| $Na_2MoO_4 \cdot 2H_2O$ | 0.75 mg |
| $ZnSO_4 \cdot 7H_2O$ | 0.24 mg |
| $Cu(NO_3)_2 \cdot 3H_2O$ | 0.04 mg |
| KOH | Adjust pH to 6.8 |

Formulas for calculating solar-to-hydrogen energy conversion efficiencies:

$$\text{Photochemical Efficiency(PE\%)} = \frac{\text{H}_2\text{ production rate} \times \text{H}_2\text{ energy content}}{\text{absorbed light energy}} \times 100\%$$

$$\text{PE\%}(\text{Ar} - R.rubrum - \text{CdS}) = \frac{0.29\left(\frac{MJ}{mol}\right) \times 101.506\left(\frac{\mu mol}{L}\right) \times 0.18(L)}{30\left(\frac{W}{m^2}\right) \times 0.002(m^2) \times 86400(s)} \times 100\%$$

$$\text{PE\%}(\text{Ar} - R.rubrum) = \frac{0.29\left(\frac{MJ}{mol}\right) \times 44.574\left(\frac{\mu mol}{L}\right) \times 0.18(L)}{30\left(\frac{W}{m^2}\right) \times 0.002(m^2) \times 86400(s)} \times 100\%$$

$$\text{PE\%}(\text{N2} - R.rubrum - \text{CdS}) = \frac{0.29\left(\frac{MJ}{mol}\right) \times 45.31\left(\frac{\mu mol}{L}\right) \times 0.18(L)}{30\left(\frac{W}{m^2}\right) \times 0.002(m^2) \times 86400(s)} \times 100\%$$

$$\text{PE\%}(\text{N2} - R.rubrum) = \frac{0.29\left(\frac{MJ}{mol}\right) \times 24.76\left(\frac{\mu mol}{L}\right) \times 0.18(L)}{30\left(\frac{W}{m^2}\right) \times 0.002(m^2) \times 86400(s)} \times 100\%$$

The standard molar enthalpy of combustion for $H_2$ is 0.29 MJ $mol^{-1}$. The native *R. rubrum* cells produced $H_2$ under Ar atmosphere for 5 days, the daily $H_2$ yield was 44.574 μmol $L^{-1}$ $OD_{600}^{-1}$. The CdS-*R. rubrum* biohybrid cells produced $H_2$ under Ar atmosphere for 5 days, the daily $H_2$ yield was 101.506 μmol $L^{-1}$ $OD_{600}^{-1}$. The native *R. rubrum* cells produced $H_2$ under $N_2$ atmosphere for 5 days, the daily $H_2$ yield was 24.76 μmol $L^{-1}$ $OD_{600}^{-1}$. The CdS-*R. rubrum* biohybrid cells produced $H_2$ under $N_2$ atmosphere for 5 days, the daily $H_2$ yield was 45.31 μmol $L^{-1}$ $OD_{600}^{-1}$. The headspace volume was 0.18 L, light illuminance was 30 $Wm^{-2}$, illumination area was 0.002 $m^2$, total illumination time was 86400 s.
